# A comprehensive workflow and its validation for simulating diffuse speckle statistics for optical blood flow measurements

**DOI:** 10.1101/2023.08.03.551830

**Authors:** Lisa Kobayashi Frisk, Manish Verma, Faruk Bešlija, Chen-Hao P. Lin, Nishighanda Patil, Sumana Chetia, Jason Trobaugh, Joseph P. Culver, Turgut Durduran

## Abstract

Diffuse optical methods including speckle contrast optical spectroscopy and tomography (SCOS and SCOT), use speckle contrast (*κ*) to measure deep blood flow. In order to design practical systems, parameters such as signal-to-noise ratio (SNR) and the effects of limited sampling of statistical quantities, should be considered. To that end, we have developed a method for simulating speckle contrast signals including effects of detector noise. The method was validated experimentally, and the simulations were used to study the effects of physical and experimental parameters on the accuracy and precision of *κ*. These results revealed that systematic detector effects resulted in decreased accuracy and precision of *κ* in the regime of low detected signals. The method can provide guidelines for the design and usage of SCOS and/or SCOT instruments.

## 1. Introduction

An accurate and often continuous assessment of microvascular, regional blood flow has many implications for diagnosis and treatment of diseases and for the study of healthy physiology. Despite continued efforts to establish practical means for measuring microvascular, regional blood flow in a non-invasive manner, this remains an important unmet need. One potential approach uses near-infrared, coherent light and the arising speckles after its diffusion [1–4].

Coherent laser light can be used to non-invasively measure local microvascular blood flow in tissue by detecting the fluctuating speckle patterns after light interaction with the tissue [5–9]. For the purposes of this manuscript, we will focus on deep-tissue, i.e. those that utilize light that penetrates up to several centimeters, measurements using photon diffusion. This is possible since near-infrared (∼650-1000 nm) light is only mildly absorbed in most tissues.

In the field of near-infrared diffuse optics, there are two common methods for determining blood flow from laser speckles. The first consists of measuring the speckle intensity autocorrelation (*g*_2_ (*τ*)) or the electric field autocorrelation (*g*_1_ (*τ*)) over a continuous range of decay times (*τ*) to derive a blood flow index [10]. Diffuse correlation spectroscopy (DCS) [10–12] and its variants [13–15] utilize this method for quantifying the speckle statistics to determine blood flow. The second common method consists of quantifying the speckle intensity statistics using a parameter called the “speckle contrast” (*κ*). Several related techniques measure *κ* to measure blood flow. These include tomographic techniques (SCOT, scDCT) for the three-dimensional mapping of blood flow from measurement of *κ* [16, 17] and techniques to calculate one or two-dimensional maps of blood flow (DSCA, SCOS, LSF, LASCA, LSCI) [2, 8, 18–20]. Of these, some techniques (LASCA and LSCI) are non-diffuse methods and therefore only measure superficial blood flow [8, 20].

Diffuse optical methods using the laser speckle contrast can achieve similar blood flow information as DCS at an overall cheaper cost per detector channel since *κ* is an integral of *g*_2_ (*τ*) over the delay times up to a longer exposure time. In other words, common scientific cameras can be utilized as “slower” detectors. This claim has been supported by experiments [3, 21, 22], simulations [23], and most recently by theoretical analyses [24].

A thorough analysis of the measurements utilizing the intensity auto-correlation of the speckle statistics, i.e. DCS, has previously been developed and tested [25–30]. Among other uses, these works have allowed the design of components (detectors, sources) and systems that target specific goals in detection precision and accuracy in DCS.

Despite the increasing prevalence in literature of the use of speckle contrast techniques, a comprehensive method for determining the effects experimental parameters have on the accuracy and precision of *κ* has not yet been developed. Accuracy in speckle contrast values, particularly in scenarios with low levels of detected light, is important to consider as the effects of detector noise can greatly influence the detected signal. Valdes et al. [2] first described this phenomenon, and subsequently developed a noise removal algorithm to reduce the effect of detector noise on the measured value of *κ*. This algorithm has been shown to be effective, however it does not correct for all detector effects, in particular shot noise.

Previous work to optimize accuracy and precision in speckle contrast measurements includes theoretical and experimental characterization of the sampling of speckles on the precision of measured *κ* [31–33], and the effect of the imaged speckle to camera pixel ratio on the accuracy of *κ* [34–36]. These earlier works did not account for the effect of experimental sources of noise, particularly detector noise, on the measured accuracy and precision of the speckle contrast signal. Recently, this gap in the existing literature was addressed by Zilpelwar et.al. [37] through a simulation method which modeled the generation and detection of decorrelating speckles including detector noise effects. The authors demonstrated that the developed model is able to simulate both the values of *κ* as well as the noise in *κ* detected using sCMOS cameras. Using this simulation, the authors investigate the effect of speckle to pixel size ratio, exposure time, and detected photon count rate on *κ* and its signal to noise ratio (SNR) for two commercially available cameras.

We have developed a separate simulation model to Zilpelwar et.al. [37], but with a similar aim of simulating the behavior of *κ* with respect to detector noise and other experimental parameters. Our model addresses details not included in Ref. [37] such as the efficacy of the detector noise correction by Valdes et.al. [2], and the behavior of *κ* in a multi-scattering regime in a semi-infinite geometry. We are specifically interested in characterizing the accuracy and precision of speckle contrast measurements taking into consideration experimentally relevant parameters such as the noise specifications of the detectors, the exposure time of the experiments, the detected photon-count rate, the measured medium, and the sampling of the detected speckles. To this end, the developed method was first verified experimentally for its ability to simulate *κ* and the noise in *κ*. After verifying the simulation method, the method was used to study the effect of accuracy and precision of *κ* in various experimental scenarios. Finally, the simulations were used to design and optimize a system capable of measuring baseline cerebral blood flow non-invasively in an adult human.

## 2. Methods

Here we focus on two dimensional detectors (*i* × *j*) with “pixels” but the results can be generalized to other standard detectors. As will be evident later on, it is more convenient to use the square of the speckle contrast (*κ*^2^) for the analysis. We assume that the *k*^2^ is derived from sampling *n* speckles that are distributed over space (*w*_*z*_) and/or over time by repeated measurements (*w*_*t*_). These *n* speckles sampled over *w*_*z*_ and/or *w*_*t*_ are used to estimate the probability distribution of the speckle intensity. From these *n* speckles, the mean intensity (*μ* (*I*)) and the variance of intensity (*σ*^2^ (*I*)) are determined.

Even in the case of ideal detectors and light sources, the calculated values are not exactly equal to the true mean and the true variance due to the effects of limited sampling. In experiments, the situation is more complex due to additional sources that contribute to the observed photon statistics such as the detector noise which further influence the measured values of mean and variance.

Therefore, these measurement effects must be accounted for in order to experimentally determine a “corrected *κ*^2^”, or the best estimate of the true value of *k*^2^. For common detectors, these corrections include a dark frame subtraction which attempts to account for the dark and read-out signal and a statistical correction attempting to estimate the shot noise as well as the dark and read-out noise variances [2].

The speckle contrast is an alternative data-type that is used to characterize the decorrelation time (*τ*_*c*_) of the intensity autocorrelation of the speckle statistics which is more commonly utilized [24, 38]. *τ*_*c*_ is in turn dependent on several aspects such as the the optical properties of the medium, the dynamics of the scatterers, the measurement geometry, the source wavelength and more. The signals that are detected in a common detector are affected by this statistical profile which in turn affects the noise statistics. Therefore, in order to simulate realistic speckle contrast signals, we need to take all this into account and incorporate the appropriate aspects of the detectors. An illustrative flowchart of the method that has been developed is shown in Figure 1 and is further detailed below.

**Figure 1.**
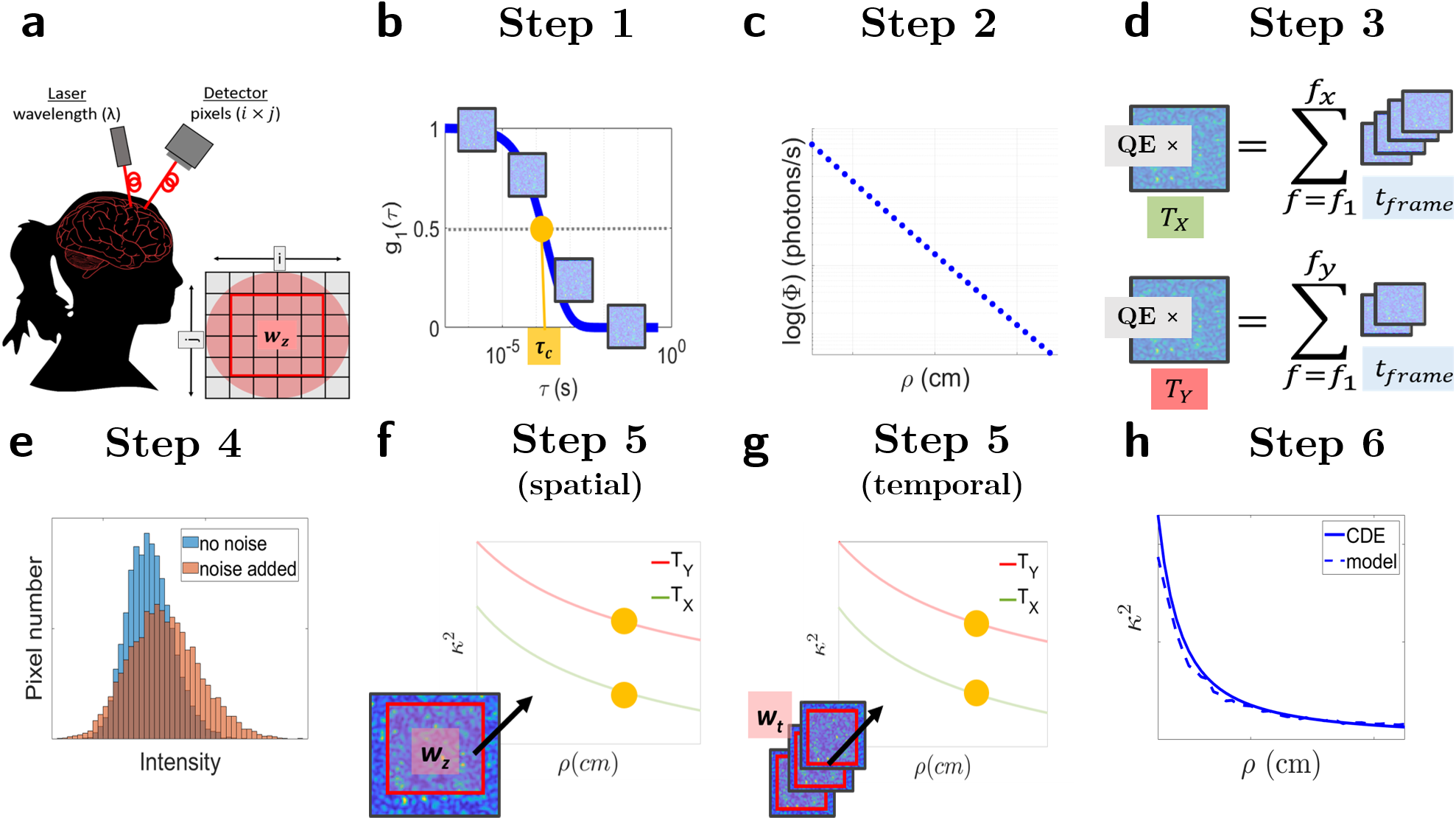
Flow chart for simulating frames of correlated speckles and *κ*^2^. These simulations aim to simulate a variety of experimental setups such as in sub-figure **a**. Depending on the experimental setup, the imaged field of view will differ. In this example, source and the detector fibers are placed a certain distance (*ρ*) from each other and are coupled to the laser and detector. The imaged field-of-view (imaged over *i* × *j* pixels includes the fiber core which in later steps will be used to calculate *κ*^2^ over a specified region of interest (*w*_*z*_). Sub-figure **b** illustrates Step 1 of the simulations. In this step, the rate at which the speckles decorrelate, *τ*_*c*_, is determined from the correlation diffusion equation (CDE). Using this value of *τ*_*c*_, consecutive frames of correlated speckles are simulated so that their electric-field autocorrelation decays with *τ*_*c*_. The intensity of these simulations are in arbitrary units, and independent of exposure time, *T*. Instead they represent speckles measured during a finite time-bin width, *t* _*f r ame*_, on the *g*_1_ curve. In order to simulate several values of *ρ*, the process illustrated in **b** can be repeated several times to simulate the *ρ* dependent change in *τ*_*c*_. In Step 2 (sub-figure **c**), the arbitrary units of the simulated frames is scaled to represent realistic values of photon current rate, Φ, in units of photons/second. In Step 3 (sub-figure **d**), an exposure time is introduced to the simulations by summing over frames. This process additionally converts the units of the simulations from photons/s to photons. Various values of *T* can be simulated from the same set of simulated frames of Step 1. In this case, the simulation of two values of exposure time, *T*_*X*_ and *T*_*Y*_, is shown. Multiplying the summed frames in units of photons by the quantum efficiency (QE) of the camera converts the units of the simulations to electrons (e^−^). In Step 4 (sub-figure **e**), the detector effects are simulated by altering the simulated intensity statistics according to the specifications of real detectors. In Step 5 (sub-figures **f** and **g**), *n* speckles are sampled over an area, *w*_*z*_ or over pixels of several repetitions of simulations to estimate a value of *k*^2^. The yellow dots represent *κ*^2^ simulated for the *τ*_*c*_ and therefore *ρ* simulated in Step 1. The two values of *T* simulated in Step 3 are also shown. In the final step (Step 6, sub-figure **h**), the discrepancies in the exact form of the speckle autocorrelation decay between the solution for the CDE for a semi-infinite medium and the developed model is corrected for.

### 2.1. The simulated experimental setup

Let us begin by detailing the canonical experimental setup that is being simulated. The exact details of the desired experimental setup to simulate may differ, however, the simulations are largely independent of these details. A visual representation of a possible setup is shown in Figure1**a**. Here, the light is delivered through an optical fiber, and detected with a separate fiber coupled to a camera. The core of the fiber is imaged with appropriate optics and all the pixels within that region-of-interest (ROI) correspond to one value of *ρ*. In a free-space system, the pixels in the imaged field of view could correspond to different values of *ρ*.

We assume that a coherent light source of wavelength *λ* is utilized. The photons, once in the medium, undergo absorption and scattering events. The probability per unit length the photons are absorbed is estimated by the absorption coefficient (*μ*_*a*_ (*λ*)). The reduced scattering coefficient 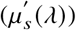 is used to estimate the total length which after a few scattering events leads to the randomization of the photon direction. In other words, after a photon traverses a distance few times the 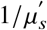, the light can be considered diffuse [39]. This diffuse light is measured at a distance *ρ* away from the source. As a rule-of-thumb, *ρ* is related to the mean probed depth by the measured light so that in order to measure deeper tissue, canonical experiments utilize longer *ρ*.

If the light source is of sufficiently narrow bandwidth (long coherence length) [40], then the so-called “diffuse laser speckles” and their statistical fluctuations can be observed. The electric-field (*g*_1_) or the intensity (*g*_2_) autocorrelation of the detected speckles are functions of parameters related to the experimental setup (e.g.*ρ* and *λ*) and the properties of the measured medium including *μ*_*a*_, 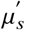, the ratio of the moving scatterers to the static ones (*α*) and the mean-squared displacement of the scatters (Δ*r*^2^). For most experiments, the “effective” particle/scatterer diffusion coefficient weighted by *α* (*αDb*) is measured as a “blood flow index” (BFI). For further details see Refs. [7, 10, 41]. The decorrelation time, *τ*_*c*_ (normally defined as the time *g*_1_ decays to 1/*e* [20]) was defined for the purpose of these simulations as the time at which *g*_1_ decayed to 0.5 and is also a function of these parameters.

### 2.2. Speckle statistics detected by a two dimensional detector array

We have simulated *k*^2^ for tissue with specific optical properties and blood flow by simulating consecutive frames of correlated speckles which simulate their electric field autocorrelation with a decorrelation time, *τ*_*c*_, defined by the solution of the CDE for a semi-infinite medium [10]. The methodology presented is independent of this solution and other solutions (layered, heterogeneous, numerical) of the CDE could be utilized. For clarity, electric-field autocorrelation curves following the solution of the CDE will be referred to as 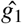, while the simulated electric-field autocorrelation curves are referred to as 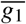. While the two are similar, there are slight differences which are discussed below. Furthermore, the theoretical value of *k*^2^ derived from the CDE will be referred to as 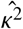 while the simulated values will be referred to as 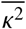.

In the first step of the simulation pipeline (Figure 1**b**), *τ*_*c*_ is derived from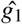. The derived value of *τ*_*c*_ was used to simulate frames of individual speckles by modifying the copula method developed in Ref. [42]. This method simulates consecutive two dimensional matrices of numbers that are correlated to each other by using a mathematical copula. Furthermore, the statistical profile of each matrix reflects the probability distribution of speckle intensity. Therefore, each individual matrix can be considered as a camera frame acquired in a speckle contrast experiment. These matrices are referred to as “frames” (*f*) simulating pixel coordinates *i, j* while imaging speckles with diameter, Ø. Ø behaves as a scaling factor to put physical units for the pixel size since the speckle diameter is approximately equal to the wavelength of light being used. Therefore, choosing Ø to be equal to three pixels for a system modeling *λ* = 785 nm will scale the width of each pixel to be equal to approximately 262 nm.

The autocorrelation, 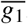, of the first frame, *f* = *f*_1_ to the k^th^ frame, *f* = *f*_*k*_ is given by

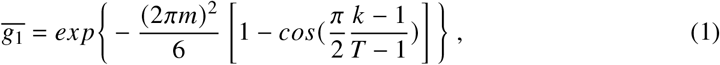

where *k* is the frame number and *m* is a parameter related to the decorrelation of the frames. In our adaptation we have defined *m* to be a function of *τ*_*c*_. Since *τ*_*c*_ has been defined as 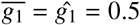 then

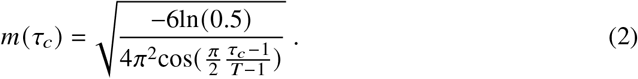

Each of the individual simulations of 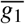 consisting of *f* = *f*_*N*_ frames of speckles patterns constitute an experiment, defined by *ϵ*. This process together with notation is illustrated in Figure 2. The basic method simulates *β*, an experimental parameter related to the coherence of the light source and the detection optics [43], equal to one. However *β* can also be simulated for other values by following the method of Ref. [42].

**Figure 2.**
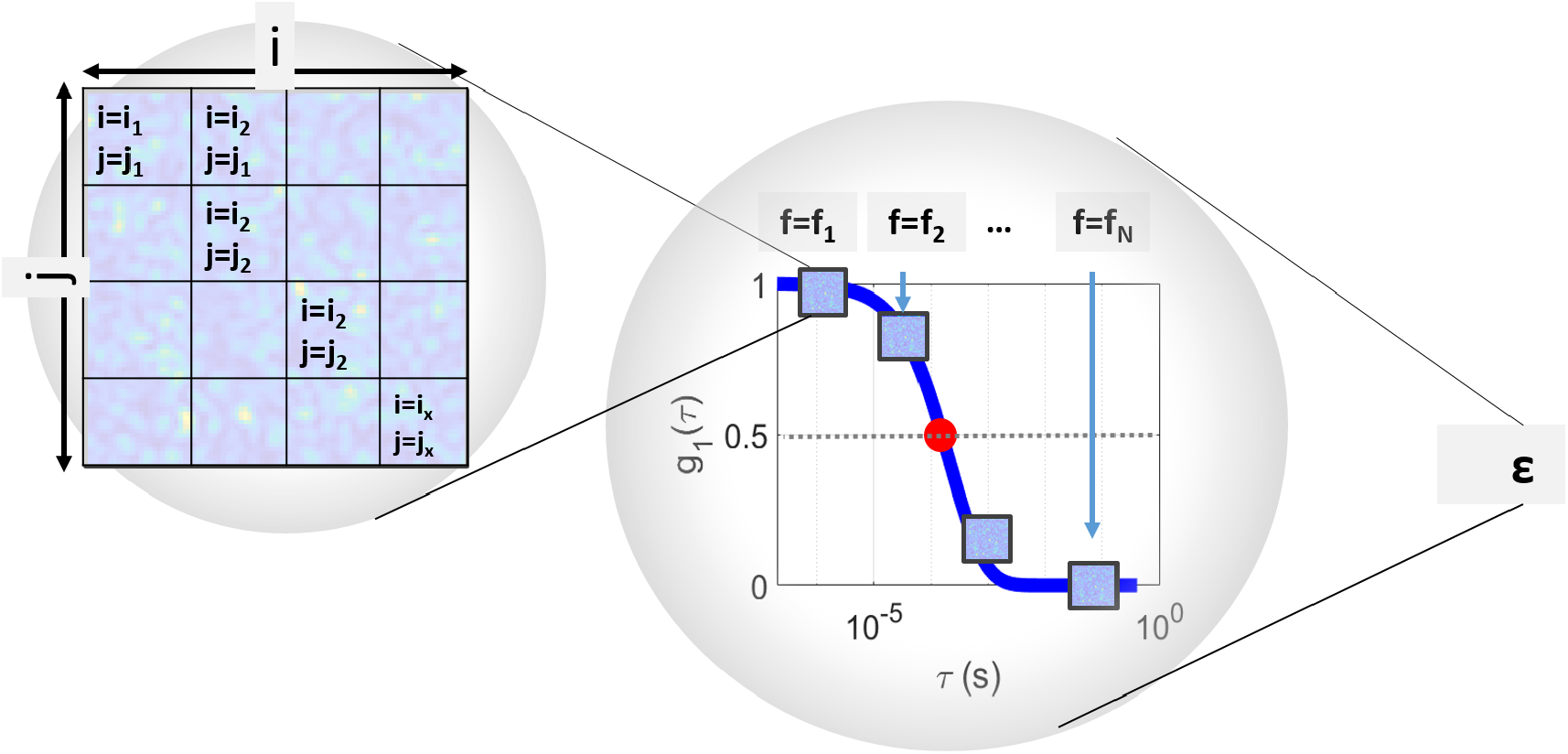
Illustration of how frames with a defined *τ*_*c*_ are simulated. First individual speckles are simulated on a grid of *i* × *j* pixels. These individual frames, *f*, are correlated to each other and their electric-field autocorrelation, 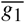, decay according to *τ*_*c*_ defined from semi-infinite theory (Figure 1). One full simulation of a theoretical *g*_1_ curve 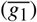 consisting of *f*_*N*_ frames corresponds to one experiment, *ϵ*. This process is repeated several times resulting in several simulations of *g*_1_.

The simulations are simulated in arbitrary copula units. In addition, the frames are only dependent on *ρ* and every simulated frame represents a point on the 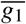 curve with a finite time-bin width, *t* _*frame*_. Since each frame has a defined *ρ* and is simulated over an array *i* × *j*, the complete notation is, 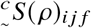. In this notation, the pre-superscript indicates the units of the simulated frame. In this case, *c* refers to the arbitrary “copula” units. The pre-subscript, ∼, indicates that no effect of detector noise has been included in the simulated frame. The indices *i, j* and *f* refer to the pixel and frame.

### 2.3. Scaling detected photon intensity

In order to convert 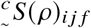 to physical units, the arbitrary copula units must be scaled to a realistic value (Fig. 1c). This is done by defining the spatial decay of light intensity theoretically or experimentally. According to the photon diffusion theory, in a semi-infinite geometry, the measured photon current rate, Φ(*ρ*), in units of photons/second, decreases with *ρ* as:

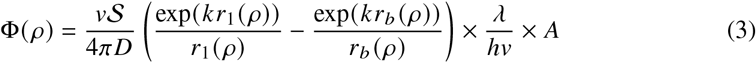

Where 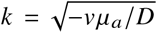, and *D* is the diffusion coefficient 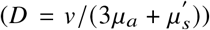, and *v* is the speed of light in medium. *r*_1_ (*ρ*) and *r*_*b*_ (*ρ*) are variables related to the boundary conditions for a semi-infinite geometry [10]. Here *h* is Plank’s constant, ***S*** is the source irradiance in units W/cm^3^, and *A* is the pixel area. It is noted that *A* in the simulations is related to the speckle size, Ø, such that *A* = *λ* Ø.

Alternatively, experimental values of Φ (*ρ*) can be used to simulate the photon current rate at the detector. In this case, the average measured photons per second at specified values of *ρ* (divided by the quantum efficieny of the specified detector) can be used to approximate the photon current rate.

Once Φ(*ρ*) has been established, whether theoretically or experimentally, the simulated frames are scaled using Φ(*ρ*) to convert them to a physically meaningful unit of photons/second, denoted as 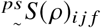. This is evalulated through the normalization of 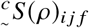 with its mean over simulated frames, 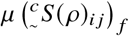:

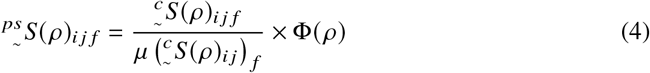

### 2.4 Introducing exposure time to the simulated frames

The next step (Fig. 1**d**) requires converting the frames of equal frame widths, *t* _*f rame*_, to frames with an exposure time, *T*_*x*_. These frames are denoted as 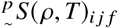 and are in units of photons.

This is done by adding *N* = *T*_*x*_/*t* _*f rame*_ consecutive frames:

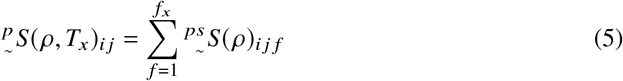

Note that with the introduction of exposure time, the simulated frames drop their indexing of *f*.

Finally, the simulated frames are converted from photons to electrons:

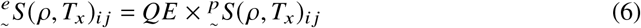

Where *QE* is the quantum efficiency of the camera.

Table 1 summarizes the introduced notation to refer to the simulated frames.

**Table 1.**
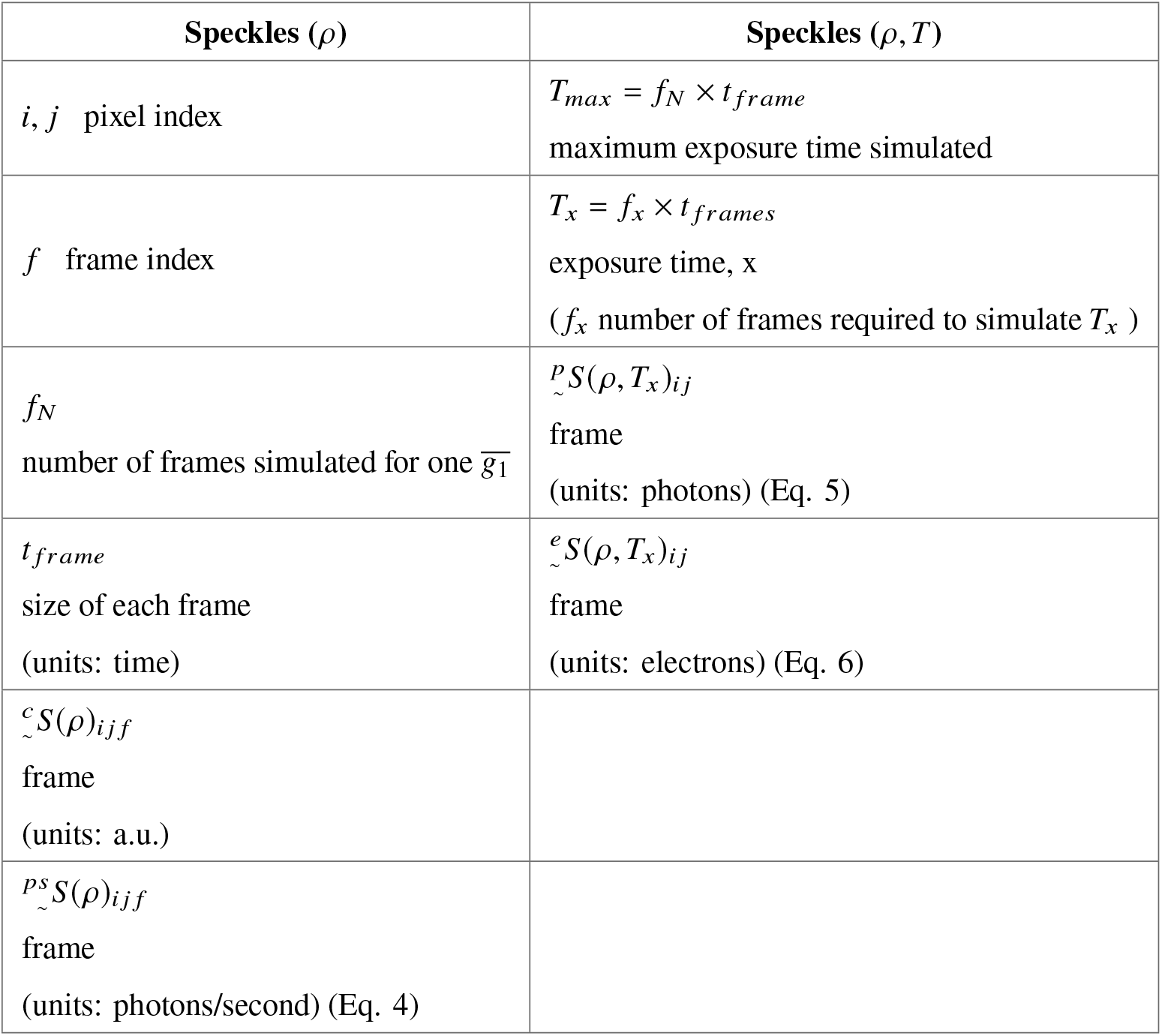
Table of definitions of the simulated speckle patterns including conversion of units from arbitrary simulation units with no *T* dependency to electron units with *T* dependency. In the notation for the simulated frames, the pre-superscript indicates the units of the simulated speckle intensities while the pre-subscript, ∼, indicates that no noise has been added.

### 2.5 Detector Noise

The final step before using the simulations to calculate 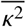 is to simulate the effects of the main types of detector noise on the simulated frames previously described, namely: photon shot noise, dark signal non-uniformity (DSNU), dark current shot noise, and read-out noise [44, 45]. This step is illustrated in Fig. 1**e**. To simulate detector noise, the distribution of each of the types of noise is considered, and random numbers are generated following the distribution. The notation used to describe the generation of random numbers and their distributions is shown in Eq. 7

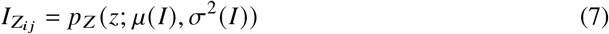

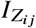 is the random number generated representing a certain intensity (in e^−^) at pixel *i,j*, 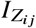 originates from a distribution, *p*_*z*_, with a mean value of intensity, *μ* (*I*), and variance, *σ*^2^ (*I*).

Photon shot noise is a Poisson distributed noise source [44, 46]. Using the notation in Eq. 7, the contribution of photon shot noise at each pixel *i, j* is described as:

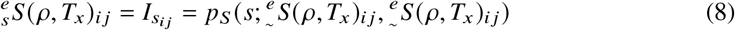

Where we have applied the definition of a Poisson distribution, *μ* (*I*) = *σ*^2^ (*I*). In this case 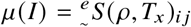 (i.e. the measured intensity in e^−^ (Eq. 6)). We have also included a new notation 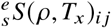. The pre-subscript, *s*, denotes the application of shot noise on the simulated frame.

DSNU and dark current noise along with read-out noise are not directly applied to 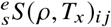, instead independent dark frames are simulated and then added to 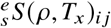.

DSNU is simulated by simulating individual pixels of logistically distributed random numbers [46]:

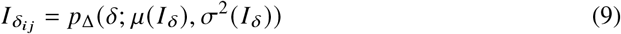

Where *μ* (*I* _*δ*_) and *σ*^2^ (*I* _*δ*_) are the mean and variance of the DSNU specific to each detector. Their values can typically be found in camera specification sheets. The variance of a logistic distribution is given by 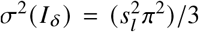 where *s*_*l*_ is the shape parameter of the logistic distribution.

The dark shot noise, similar to the photon shot noise (Eq. 8) is simulated by applying Poisson distributed random numbers [44] to each pixel simulated in Eq 9:

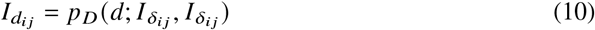

Finally, read out noise is simulated by assuming that it is a normally distributed noise source [47]. Read out noise in CMOS cameras is added at each pixel and is independent of the dark noise and the detected signal. Therefore, the contribution of the read out signal at each pixel, 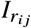, is simulated:

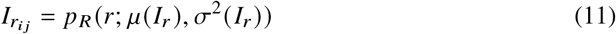

where the mean and variance of the read-out signal (*μ* (*I*_*r*_) and *σ*^2^ (*I*_*r*_)) are specific to each detector and can be found in specification sheets or estimated from online camera simulators.

The total dark frame, *df*, is then given by

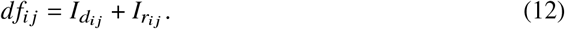

Putting everything together, the frames with shot noise, DSNU, dark shot noise, and read-out noise, 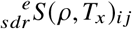, are given by:

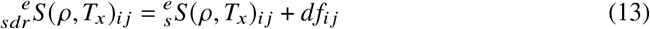

To generalize the notation, the pre-subscript *N* indicates a general noise source. In other words, 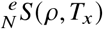 is shorthand for speckle intensity frames in units of electrons with unspecified noise, *N*, added. *N* can take values:

- ∼ : no noise
- *s* : shot noise added
- *sdr* : shot noise and dark frame added (dark and read out noise)
- *sd*^′^*r*^′^ : shot noise and dark frame added, dark frame offset subtracted (dark and read out noise corrected)
- *s*^′^*d*^′^*r*^′^: shot noise and dark frame added, dark frame and shot noise corrected.

The definitions and notation for simulating detector noise is summarized in Table 2:

**Table 2.**
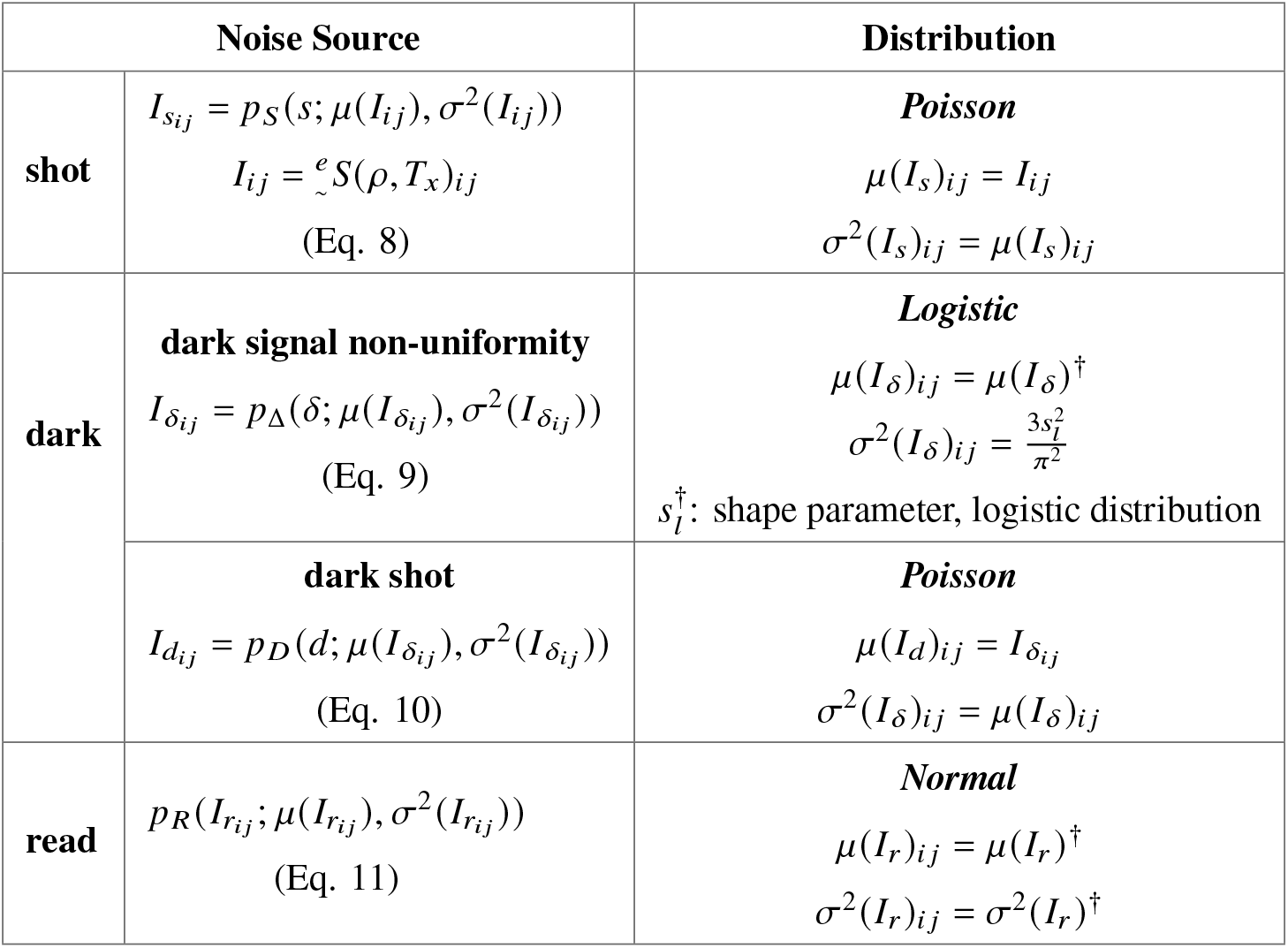
Table of definitions of the noise sources that are included in the simulations along with their corresponding distributions. The notation *p*_*z*_ (*z*; *μ, σ*^2^) is used to define random numbers, *z*, originating from a distribution, *p*_*z*_, with a mean value of, *μ*, and variance, *σ*^2. †^denotes parameters that can be found in camera specification sheets.

### 2.6 Speckle Contrast

The final steps of the simulation pipeline require the calculation of 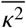 using the frames that have been simulated. In the first step, 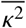 is directly calculated using the simulated frames. The calculation of 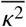, as in a real experimental setting, can be done temporally or spatially depending on how speckles are sampled. Independent of the domain in which 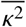 is simulated, it should be noted that since the speckle decorrelation was modelled as a single exponential (Eq. 1), the physically more realistic semi-infinite model of the speckle decorrelation follows a double exponential model [10]. A correction was applied in order to simulate a model corrected value of 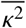 denoted as 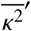. Previous work in developing a successful DCS noise model also applied a single exponential model in order to model noise [25, 48]. Therefore, while the value of 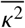 will be affected by the model used for 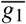, the noise is well described using the simplified single exponential model. The definitions and notation related to 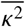 are summarized in Table 3. The following sections will describe their calculations.

**Table 3.** Table of definitions for *κ*^2^. Three different variations of *κ*^2^ are calculated: first *κ*^2^ calculated directly from the integration of the double exponential *g*_1_ from CDE. This is 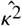. Secondly, *κ*^2^ calculated directly from the simulated frames whose 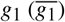 follows a single exponential form. This is 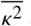 and outlined in Section 2.7. Thirdly, the model differences due to the differences in *g*_1_ is corrected. This is 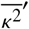 and is outlined in Section 2.8. Moreover, 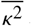 and 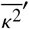 can be calculated either spatially or temporally.

| $\kappa^2$ | Spatial $\kappa^2$ | Temporal $\kappa^2$ |
| --- | --- | --- |
| $\hat{g}_1$<br>electric-field autocorrelation curve<br>CDE, semi-infinite solution [10] | $w_z = [i_\zeta j_\zeta, i_\xi j_\xi]$<br>“spatial window” of pixel area | $w_t = [\epsilon_\zeta, \epsilon_\xi]$<br>“temporal window” of experiments |
| $\hat{\kappa}^2$<br>**derived from $\hat{g}_1$** | $\mu(I_\epsilon)_{w_z}$<br>mean intensity over $w_z$ | $\mu(I_{ij})_{w_t}$<br>mean intensity over $w_t$ |
| $\overline{g}_1$<br>simulated autocorrelation curve<br>(Eq. 1) | $\sigma^2(I_\epsilon)_{w_z}$<br>variance of intensity over $w_z$ | $\sigma^2(I_{ij})_{w_t}$<br>variance of intensity over $w_t$ |
| $\overline{\kappa^2}$<br>**derived from $\overline{g}_1$** | $\overline{N\kappa^2}_\epsilon = \frac{\sigma^2(I_\epsilon)_{w_z}}{\mu^2(I_\epsilon)_{w_z}}$<br>**spatial $\kappa^2$ (Eq. 14)** | $\overline{N\kappa^2}_{ij} = \frac{\sigma^2(I_{ij})_{w_t}}{\mu^2(I_{ij})_{w_t}}$<br>**temporal $\kappa^2$ (Eq. 15)** |
| $N\gamma = \overline{\kappa^2} - \overline{N\kappa^2}$<br>bias term (Eq. 19) | | |
| $\overline{N\kappa^2}' = p_K(k; \hat{\kappa}^2 + N\gamma, \sigma^2(\overline{N\kappa^2}))$<br>**corrected for semi-infinite theory**<br>(Eq. 20) | | |

### 2.7 Model uncorrected speckle contrast

So far the process for simulating the detection of speckle statistics on a 2D detector array and the detector properties (Fig. 1 **b** to **e**) has been described. These steps can be repeated in order to simulate several experiments (*ϵ*, Fig. 2) for several different values of *τ*_*c*_ and therefore *ρ*, for calculating 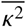 in the temporal domain over *w*_*t*_, or for determining 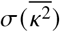.

The next step in the pipeline is to use these frames to calculate values of 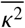 (Fig. 1 **f** and **g**).

As mentioned previously, 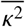 can be measured spatially or temporally i.e. speckle statistics can be determined spatially by using an area, *w*_*z*_, of pixels or temporally over the pixels in a set of experiments, *w*_*t*_.

Spatial 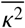 is given by:

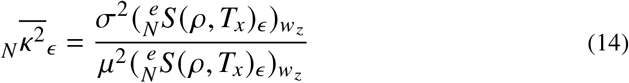

Where 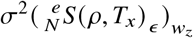 is the variance of the speckles and 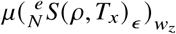 is the mean of the speckles, both calculated over the window *w*_*z*_ for each experiment, *ϵ*.

Similarly, temporal 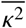 is given by:

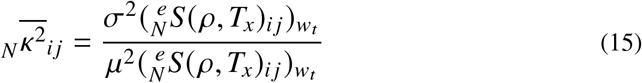

Where in this case, the variance and means of the speckle intensities are calculated over a temporal window of many experiments *w*_*t*_ for a set of *i* × *j* pixels.

With 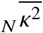 simulated, noise correction must be applied. To do this, the noise correction method outlined in [2] was used. Here we outline the correction for spatial 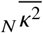, but the same principles apply for temporal measurements.

Briefly, in order to correct for the dark and read signal offset in 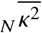, a new dark frame, *df*_*corr*_, is simulated using Eq. 12. The new dark and read signal offset corrected speckles frames is given by:

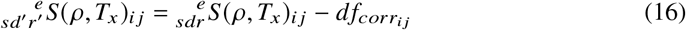

After the dark frame offset is corrected, the additional variance due to shot 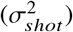 and the dark frame (dark and read out noise, 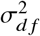) is corrected by subtracting their respective variances from the signal variance, 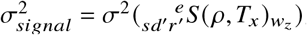.

Putting everything together, the shot, dark, and read noise corrected value of 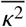, i.e. 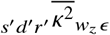, is given by:

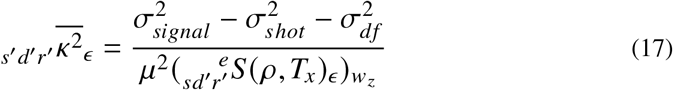

Where 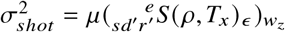 and 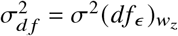.

Variations in the noise correction can also be simulated. For example, the shot noise only added frames, 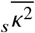, can be corrected in the following way:

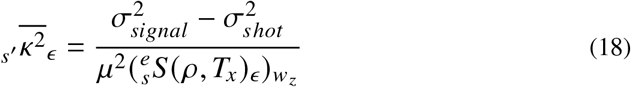

Where in this case, 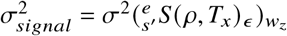 and 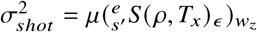.

### 2.8 Model corrected speckle contrast

In these simulations, two forms of the electric field autocorrelation function have been introduced: 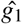 and 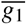, and crucially the decorrelation of the latter was modeled from the decorrelation time of the former. However, the two are described by two different exponential functions meaning that the values of *k*^2^ derived from the two will differ. In particular, 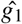 describes a measurement in a semi-infinite medium and a multi-scattering (diffuse) regime. Since 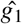 is a more realistic solution to the CDE, rather than working with 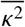 derived from 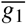, we introduce another variable, 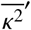, which is the model-corrected value of 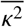.

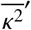 is derived from both 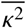 and 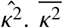 values are used to simulate the offset or bias (*γ*) in *k*^2^ due to noise, as well as to simulate the expected variance of *κ*^2^ over *ϵ*. The CDE solution of 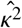 is then used to scale the value of 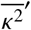 to the expected value of speckle contrast when measuring in a semi-infinite geometry.

The bias term, *γ* is defined as:

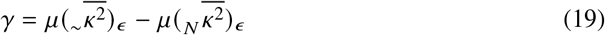

Finally 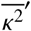 values are generated by generating normally distributed random numbers, *k*, with mean equal to 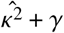 and variance equal to 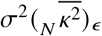 :

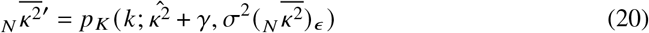

### 2.9. Using the simulations to evaluate system performance

A primary motivation for developing a speckle contrast model is to evaluate the performance of such systems. Performance of simulated systems has been evaluated by its accuracy and precision. In this context, accuracy refers to the percent error of 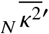 from its CDE solution, 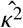, and was defined as 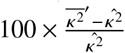. Precision is a measure of how variable a repeated measurement is and has been evaluated by its coefficient of variation (CV) as a percentage defined as the ratio of standard deviation of repeated experiments of 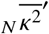 to its mean: 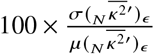. Maximum accuracy and maximum precision correspond to the minimum values in these metrics.

### 2.10. Experimental setup (A) to validate simulations

The speckle contrast noise model was validated by comparing experimental results to the simulated noise for a range of exposure times. A multi-mode fiber delivered light (785nm, Crystalaser, Reno NV, USA), onto a liquid phantom of water, intralipid and ink. The resulting speckle pattern was imaged onto an sCMOS camera (Orca Fusion-C14440-20UP, Hamamatsu Photonics K.K., Hamamatsu, Japan) using a multi-mode fiber (910 *μ*m core, 0.22 NA) and objective lens (f = 11 mm). The value of *β* was measured to be approximately 0.2, and Ø was adjusted to be approximately 4 pixels.

*τ*_*c*_ of the system was obtained by simultaneous recording *g*_2_ of the system using a single mode fiber coupled to a standard DCS device. The detector fibers of both the SCOS system as well as the DCS system were placed at a distance *ρ* = 0.8 cm from the source. The performance of the simulations was compared to the experimental results by evaluating the standard deviations of 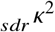 of both over 100 experiments. In addition, the expected signal-to-noise-ratio (SNR) was also evaluated considering 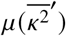 to be equal to the average value of 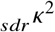 over 100 experiments (Eq. 20). SNR is defined as the ratio of the average value of the signal over the noise. The experimental values of 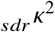 was calculated over a horizontal row of 1032 pixels. The simulated SNR was defined as the ratio of the standard deviation of the experimentally obtained values of 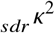 to the average value of 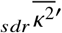 over 100 simulated experiments, *ϵ*, calculated over 1032 simulated pixels.

### 2.11. Experimental setup (B) to optimize and design a speckle contrast system

The speckle contrast noise model was further used to design a speckle contrast system and define the required detected electron count rate (e^−^/pixel/second) in order to accurately measure blood flow in the adult human brain. An sCMOS camera by Basler (daA1920-160um, Basler AG, Ahrensburg, Germany) was considered and simulated due its lightweight (15 g), compact size (19.9 mm x 29.3 mm x 29 mm) and cheap price (<300 €). Measurements were chosen to be taken at *ρ* of 2.5 cm and *T* of 5 ms.

The required detected electron count rate to accurately measure *k*^2^ was determined by attenuating a 785 nm laser (Crystalaser, Reno NV, USA) on a liquid phantom using a fiber attenuator (OZ Optics, Ottawa Ontario, Canada). The diffuse light was imaged onto the camera using an 800 *μ*m core multi-mode fiber (0.22 NA). The imaged speckles had a size of Ø = 5 pixels. The value of *β* of the system was previously determined to be approximately 0.2. Speckle contrast data was acquired over 600 frames, and data was analyzed using an ROI of approximately 1100 pixels.

As in the setup (A) to validate the simulations, *τ*_*c*_ of the simulations was obtained from *g*_2_ recorded using a standard DCS device. In order to approximate the required detected electron count-rate (e^−^/pixel/second), a liquid phantom was prepared to have optical properties of *μ*_*a*_ = 0.1 cm^−1^ and 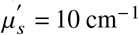. The true value of *κ*^2^ was considered to be the value of *k*^2^ measured with the highest detected intensity count rate, *I*_*max*_. Percent error of *k*^2^ as a function of the attenuated detected intensity count rates, *I*_*act*_, was therefore calculated as: 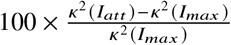.

## 3. Results

### 3.1 Verification with experimental data

The results of the simulation model were compared to experimental data of an Orca Fusion camera using the experimental set-up in Section 2.10. Details of the camera parameters are summarized in Table 4. The simulations used *τ*_*c*_ obtained from the *g*_1_ curve recorded using DCS (Figure 3 **a**). *β* was simulated to be 0.2 and Ø was set to 4 pixels to agree with the values of *β* and Ø of the experimental data. Both experimental and simulation results were obtained for exposure times ranging between 0.1 ms and 5 ms in order to cover a range of detected electron intensities. It was ensured that the average value of the simulated detected electron intensity matched the experimental data (Figure 3 **b**). The resulting experimental and simulated standard deviation of 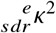 is shown in Figure 3 **c**. The calculated signal to noise ratio of *k*^2^ in Figure 3 **d**, shows good agreement of the simulations with the experimental results.

**Table 4.** Simulation parameters used to verify simulations with experimental data acquired using an sCMOS camera (Orca Fusion-C14440-20UP, Hamamatsu Photonics K.K.)

| Tissue Parameters | Detector Parameters | Speckle Parameters |
| --- | --- | --- |
| $\tau_c$: $4.18 \times 10^{-5} \text{ s}$ | QE: scaled from measurements | ∅: 4 pixels |
| | $\mu(I_\delta)$ : $0.0025 \text{ e}^-$ | $\epsilon_N$ : 100 |
| | $\sigma^2(I_\delta)$ : $0.16 \text{ e}^-$ | $w_z$ : $[0, 0; 32, 32]$ |
| | $\mu(I_r)$ : $0.93 \text{ e}^-$ | |
| | $\sigma^2(I_r)$ : $0.24 \text{ e}^-$ | |

**Figure 3.**
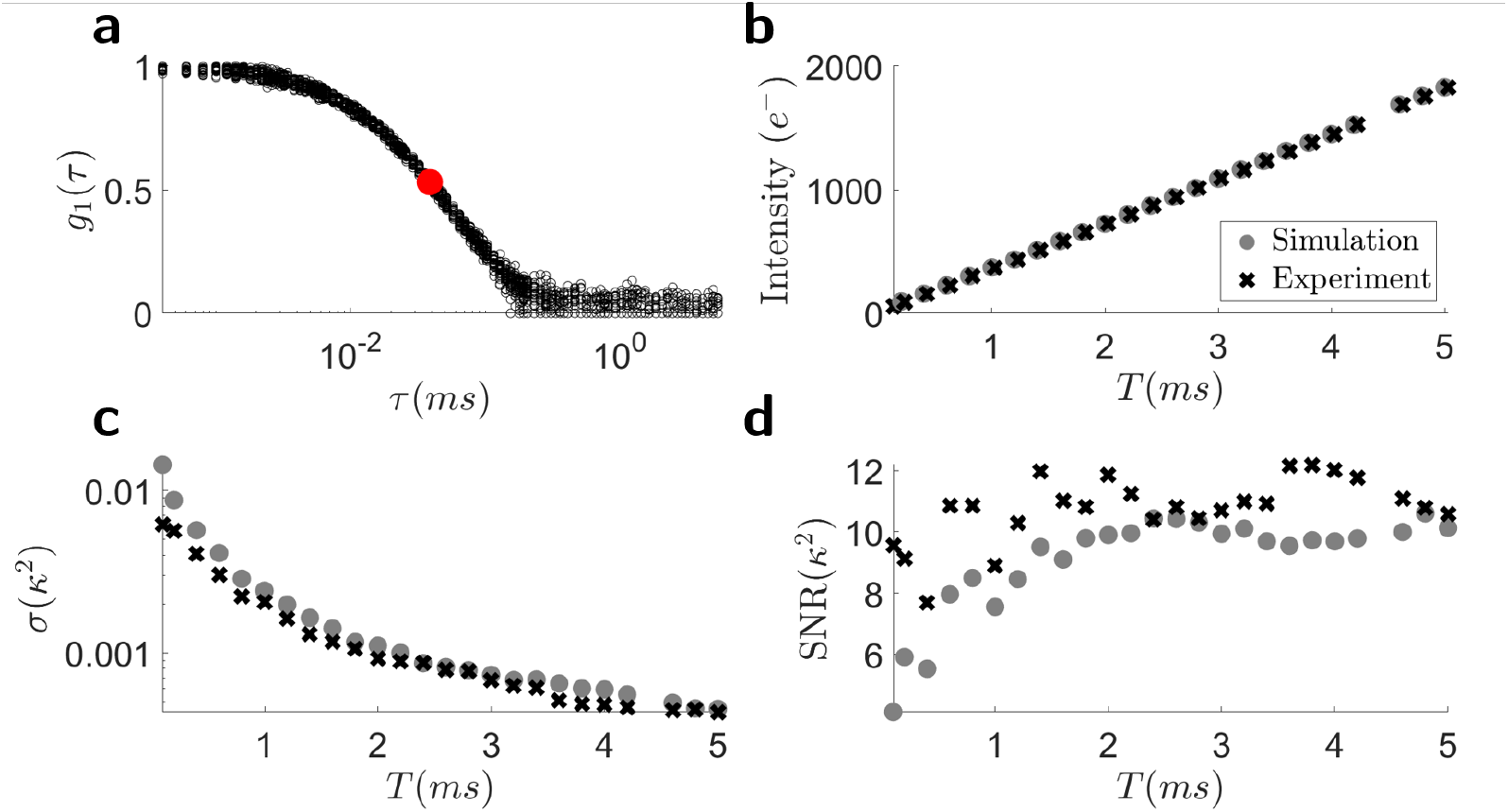
Comparison of the developed speckle contrast noise simulation model with experimental values. The number of experiments as well as the number of speckles used to obtain *k*^2^ were the same for experiments and simulations. **a**) Experimental *g*_1_ curves measured with a DCS system from which *τ*_*c*_ used in the simulations was determined (red). **b**) Average detected electrons over 1032 pixels and 100 experiments (black) and 100 simulations over 1000 pixels (grey). **c**) The standard deviation in 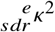 calculated by simulation (grey) and the experimental results (black). **d**) SNR from experiment (black) and simulation (grey).

### 3.2 Simulation study

Using the simulation pipeline described, we simulate speckle patterns with realistic detector noise. All simulations considered hardware consisting of a 785 nm unpolarized laser (*β* = 0.5) and a 100 × 100 pixel array detector with noise properties derived from an Orca Flash4.0 v3 CMOS camera [49]. Since the variance of read-out noise is typically not defined in specification sheets, an online simulation tool was used to approximate the value of *σ*^2^ (*I*_*r*_) [50]. Tissue with optical properties listed in Table 5 were simulated. These values were chosen as they are roughly the expected values when measuring in human tissue. 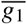 was simulated for *ρ* ranging from 0.5 to 4.5 cm for *T*_*max*_ = 5 ms. Ø was chosen to equal three pixels in order to meet the requirements of the Nyquist criteria [35, 51]. The details of the parameters used in the simulation are summarized in the table below:

**Table 5.** Parameters that were used to simulate synthetic speckles. Optical properties were chosen to mimic biological tissue, and detector parameters are based off of the properties of the Orca Flash4.0 v3 CMOS camera by Hamamatsu K.K.

| Tissue Parameters | Detector Parameters | Speckle Parameters |
| --- | --- | --- |
| $\mu_a : 0.1 \text{ cm}^{-1}$ | QE: 54.2% | ∅: 3 pixels |
| $\mu'_s : 10 \text{ cm}^{-1}$ | $\mu(I_\delta) : 0.06 e^- / s$ | $\epsilon_N : 100$ |
| $n : 1.33$ | $\sigma^2(I_\delta) : 0.16 e^-$ | $w_z : [0, 0; 100, 100]$ |
| $Db : 1 \times 10^{-8} \text{ cm}^2 / s$ | $\mu(I_r) : 2.9 e^-$ | |
| | $\sigma^2(I_r) : 0.1 e^-$ | |

### 3.3. Part I: Simulating 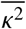

The simulated values of the decorrelation time, *τ*_*c*_, as a function of source-detector separation, *ρ*, is shown in Fig. 4 **a**. As expected from theory, the speckle autocorrelation decays faster with increasing *ρ* [10], confirming that the modified copula method for simulating decorrelating speckle intensity replicates the expected dynamics from theory. In Fig. 4 **b**, 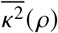 calculated by integrating the simulated speckle electric field decorrelation curves, 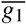 (Eq. 1) for three different exposure times is shown. As expected from theory, 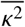 decreases with increasing *ρ* and increasing *T*.

**Figure 4.**
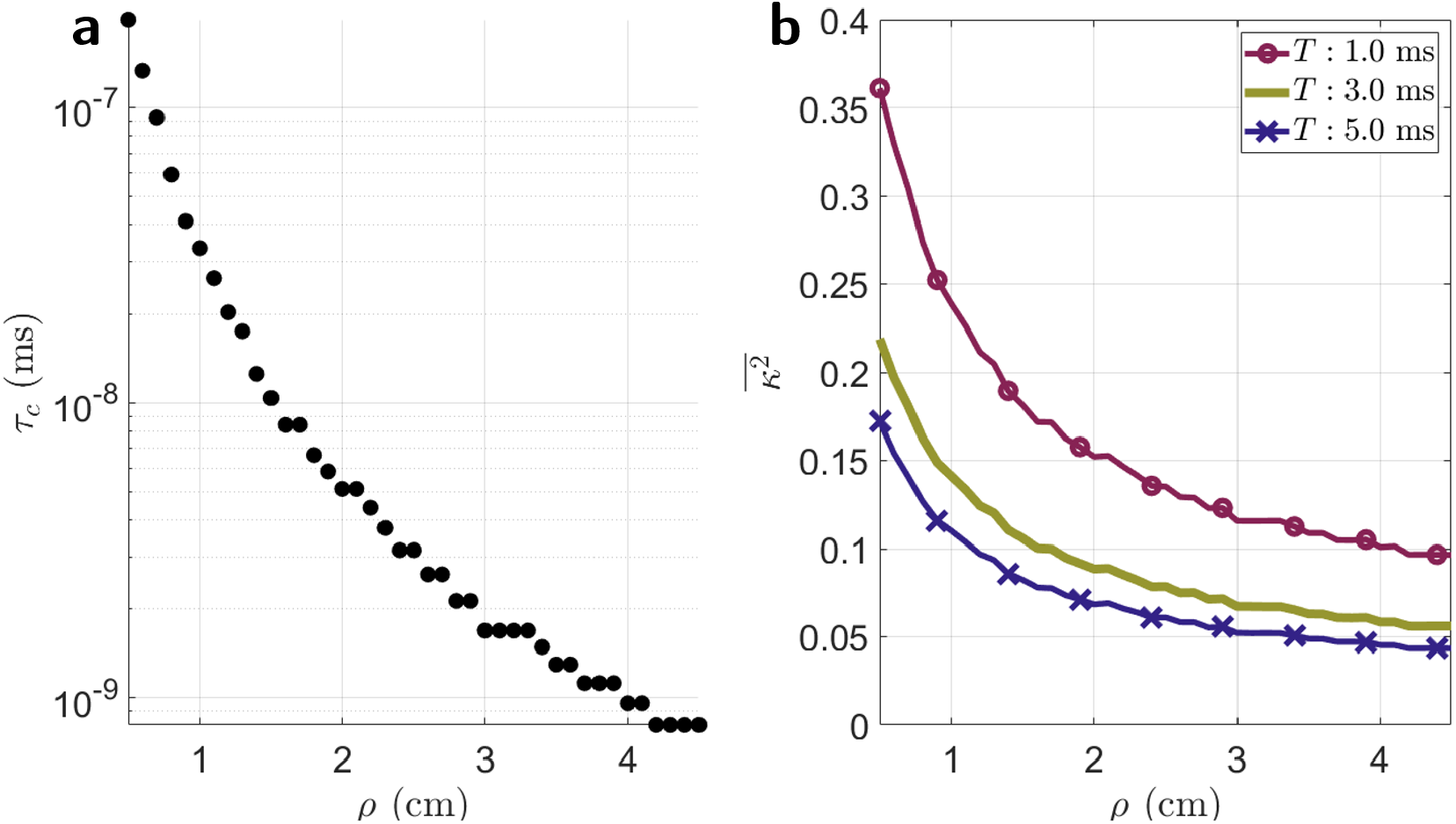
**a**) simulated values of *τ*_*c*_ in ms. A clear decrease in *τ*_*c*_ with increasing *ρ* is seen. **b**) 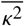 at three different exposure times calculated from integrating the autocorrelation, 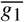, of the simulated speckles.

The simulated detected number of electrons 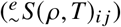 for different *ρ* at two different *T* for all 100 simulated experiments are shown in Fig. 5 **a** and **d**. Including detector effects in the simulations results in deviations of the average value and variance from the ideal detected electron intensity value. This effect is *ρ* and *T* dependent. For all values of *ρ* and *T*, the average value of the electron intensity does not deviate from the ideal case when only shot noise is simulated (N: *s*). However, in the regime of lower detected electron counts originating from speckle signal, i.e. at longer *ρ* and shorter *T*, there is an increased variance in the shot noise included detected electron intensity. Furthermore, at short *T*, it is seen that the addition of a dark frame (N: *sdr*) visibly leads to a deviation in the average value of the detected electron intensity at *ρ* = 2 cm, while the same deviation for higher *T* is not observed until approximately *ρ* = 4 cm. This is explained by the properties of the camera that were simulated. In this case, the dark current, a *T* dependent signal, was significantly smaller than the read out signal, a *T* independent signal, for the exposure times shown (*μ* (*I*_*d*_) = 6 × 10^−6^ *e*^−^ and *μ* (*I*_*d*_) = 3 × 10^−4^ *e*^−^ for *T* = 0.1 ms and *T* = 5 ms respectively, compared to *μ* (*I*_*r*_) = 2.5*e*^−^). Therefore, while dark noise is a *T* dependent noise source, the effect of adding a dark frame appears more significant at shorter *T* due to the high read-out signal relative to the speckle signal. Subtracting a dark frame (N: *sd*^′^*r*^′^) corrects this deviation. However a dark frame subtraction does not correct the increase in variance of the detected signal due to shot, dark, and read-out noise terms.

**Figure 5.**
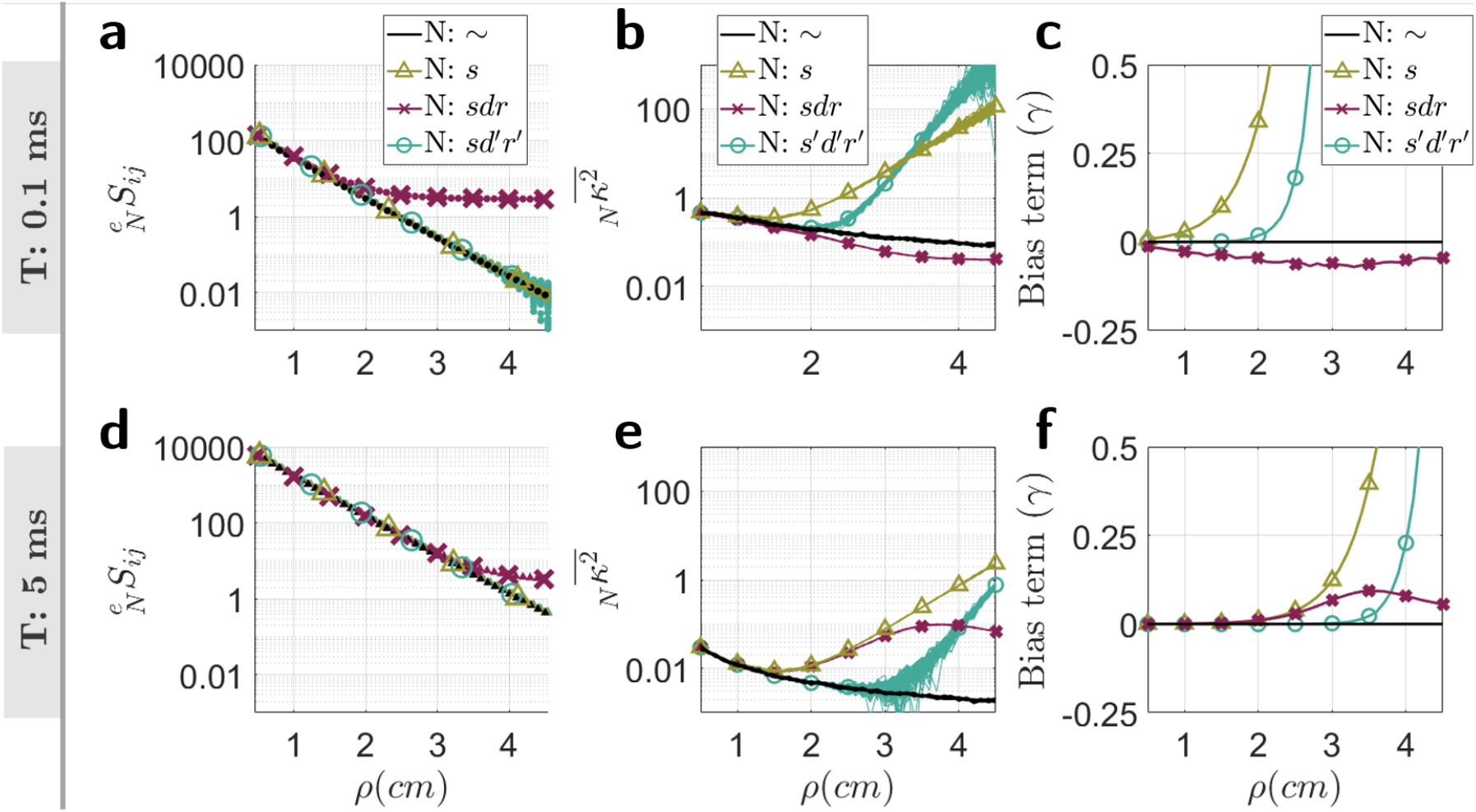
Simulation of 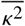 from the frames of synthetic speckles. **a, d**) Φ (*ρ*) for two different exposure times (*T* = 0.1 ms and *T* = 5.0 ms on the top and bottom rows respectively) for when no noise source are added are shown as well as for when noise sources are added and when a dark frame is subtracted. **b, e**), the values of 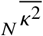 for all 100 simulated experiments. **c, f**) In order to correct for differences in theory of *g*_1_ between the double exponential form of the semi-infinite model from CDE and the single exponential copula model, a bias term *γ* is calculated (Eq. 19). These are shown for different variations of added noise, N, at the two simulated exposure times.

These observations are carried through to Figure 5 **b** and **e** where the values of 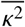 are plotted. At shorter *ρ* and for both values of *T*, simulation of detector effects show very little deviation from the ideal, no detector noise added case. However, with increasing *ρ*, there is a noticeable deviation, as expected from experiments [2]. In the case of addition of shot, dark, and read-out noise (N: *sdr*), it is seen that for *T* = 0.1 ms (Figure 5 **b**), 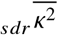 begins to deviate from the ideal case, at approximately *ρ*=2.0 cm. At *T* = 5.0 ms (Figure 5 **e**), 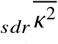 begins to deviate from the ideal case from approximately *ρ*=1.5 cm. Correcting for detector effects by applying a dark frame subtraction and correcting for shot, dark, and read-out noises (N: *s*^′^*d*^′^*r*^′^) results in a larger range of *ρ* for which 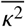 agrees with the ideal case for *T*=5.0 ms, to about *ρ*=3 cm. However, the same correction does not obviously perform as well for *T*=0.1 ms (Figure 5 **b**), with detector effects correction (N: *s*^′^*d*^′^*r*^′^) apparently performing worse than the uncorrected case (N: *sdr*). This last observation should not be interpreted as a failure in the correction of noise, rather it is a reflection of the origin of the electron signal in this regime. Referring back to the plot of the detected intensity (Figure 5 **a**), at *T*=0.1 ms, the majority of the detected electron signal after *ρ*=2 cm originate from the detector rather than from speckles. Therefore, without applying corrections, any value of *k*^2^ in this regime is not a reflection of speckle contrast, rather reflects a “detector signal″ contrast.

The bias term, *γ* (Eq. 19), is shown in Fig. 5 **c** and **f** and reflects the offset of 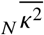 from the no noise added case, 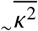. These were used to calculate the average theory corrected value of *k*^2^ with simulated detector effects 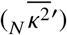. For the remaining results, only the case of N = *s*^′^*d*^′^*r*^′^ will be considered as this is the case of most interest in any experiment. The theory corrected values of *k*^2^ are shown in Fig. 6 **a** and **d**.

**Figure 6.**
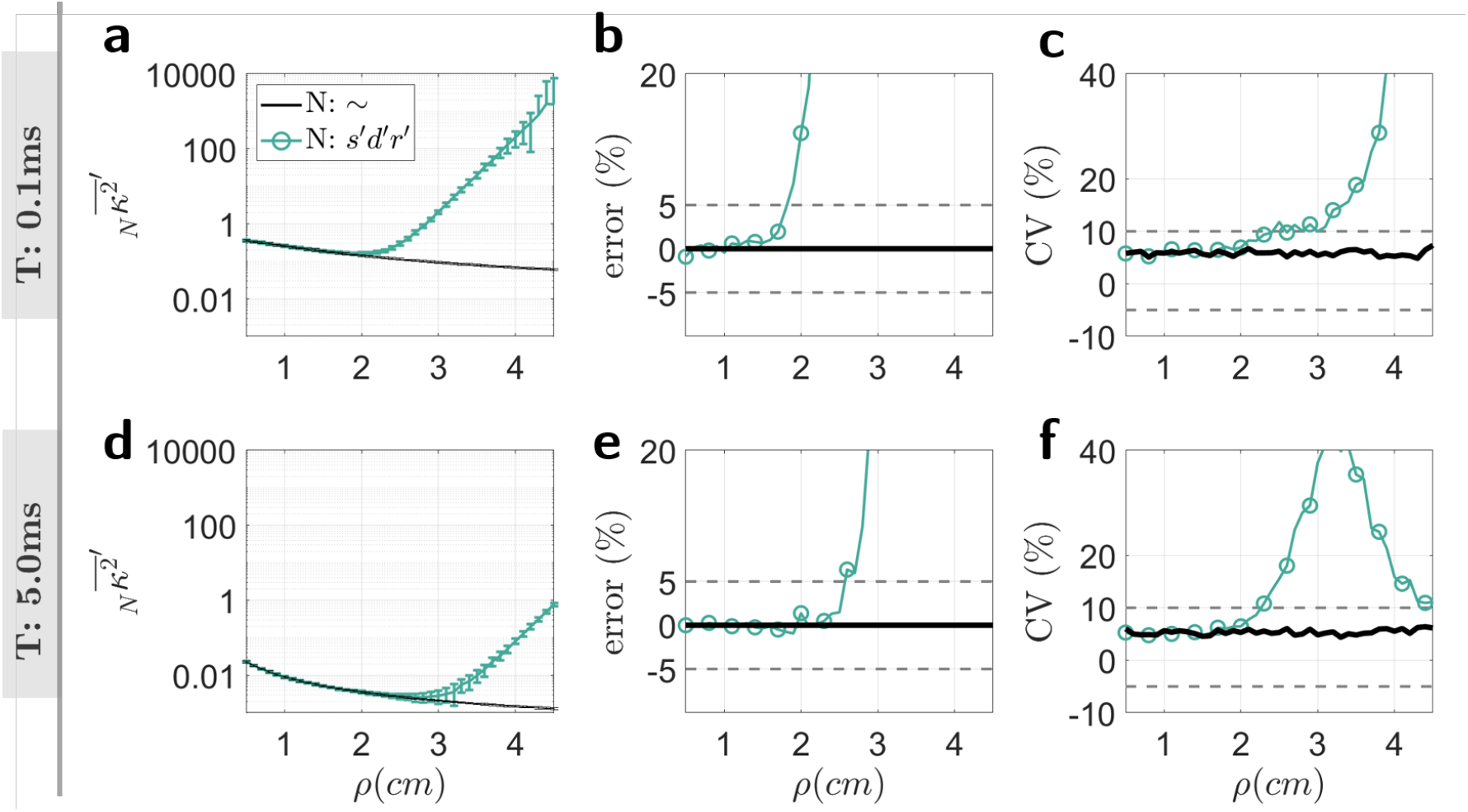
**a, d**) Simulation of theory corrected values of speckle contrast, 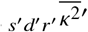. **b, e**) Accuracy (percent error) of 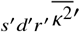. **c, f**) Precision (coefficient of variation) of 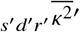

Theory corrected values of speckle contrast, 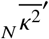, were calculated from Eq. 20. The final averaged value of the simulated 500 normally distributed random values of 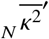 for *T* = 0.1 ms and *T* = 5 ms are plotted in Fig. 6 **a** and **d**. Error bars reflect the standard deviation. The accuracy of 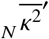 is shown in Fig. 6 **b** and **e**, reflected as the percent error. The percent error increases (accuracy decreases) with increasing *ρ* reaching 5% at approximately 1.8 cm for short *T* (Fig. 6 **b**) and 2.5 cm for long *T* ((Fig. 6 **e**). Similarly, the precision of 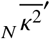, represented as the coefficient of variation (CV) also decreases (CV increases) with increasing *ρ* (Fig. 6 **c** and **d** for *T* = 0.1 and *T* = 5.0 ms respectively).

### 3.4. Part II: Using the simulations to study precision and accuracy

As seen in the previous section, effects of detector noise lead to decreases in accuracy of 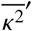 particularly in the regimes of long *ρ* and short *T*. In the next part of this analysis, the simulations are used to understand how various parameters can be changed in order to increase the usable range of *ρ* and *T* considering both precision and accuracy. In order to quantify the requirements of a SCOS or SCOT system, it is assumed that the required accuracy is within a 5% error and precision within a 10% coefficient of variation (CV) at *ρ*=4 cm and *T*=5 ms. These values were chosen for deep tissue measurement: *ρ*=4 cm corresponds to an approximate measurement depth of 2 cm. Although *ρ*=2.5 cm is considered sufficient for measuring the cortical surface going to further distances offers greath depth sensitivity and distances of between 3.0 - 4.0 cm have been used for tomographic reconstruction of human functional activation [52, 53]. *T*=5 ms was chosen in order to be able to sample at fast enough acquisition rates while also maximizing the number of detected photons (Figure 5 **d**).

In speckle contrast optical tomography (SCOT) or speckle contrast diffuse correlation tomography (scDCT) [16, 17], several source and detector positions are used in order to reconstruct a three dimensional image of blood flow. In a system incorporating nine source positions as in [54], using *T*=5 ms, this will correspond to a full acquisition rate of 22.2 Hz for *k*^2^ measured at each source position. Furthermore, 5% accuracy and 10% precision have been chosen as our targets since a 10% blood flow change corresponds to approximately 10% change in *k*^2^. A 10% change in flow is similar to what is measured in functional studies [21].

It is known that a contributing factor to the precision of *k*^2^ is the number of speckles used to determine *μ* and *σ*^2^ [31, 35]. In the previous simulations of 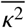, *w*_*z*_ = 100 × 100 pixels corresponding to the sampling of 1100 independent speckles. In Figure 7, *w*_*z*_ was changed to simulate the effects of the number of independently sampled speckles on the CV of 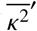.

**Figure 7.**
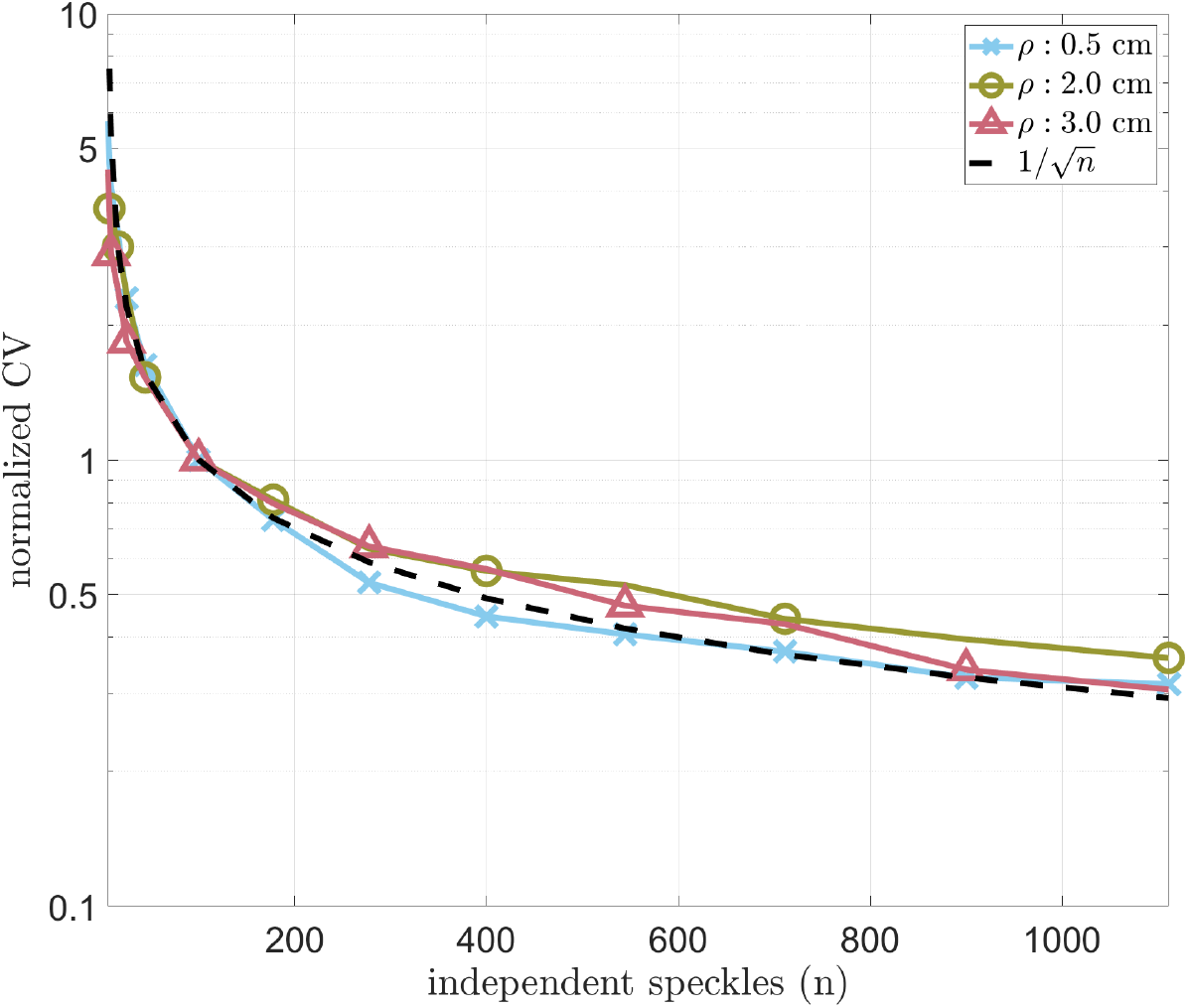
The effect of the number sampled speckles on the measured precision of 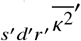 at three values of *ρ*, and *T* = 5 ms. Increasing the number of sampled speckles results in a decrease in the CV of 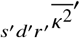.

As expected in Fig. 7, increasing the number of speckles used to calculate *k*^2^ results in an increase in the precision of *k*^2^. The decay in CV with increasing speckle number follows a square root dependency, in accordance to the theory [31]. Therefore, if the objective is to measure *k*^2^ with 10% precision at *ρ*=4 cm and *T*=5 ms, *w*_*z*_ must be increased from 100 x 100 to approximately 170 x 170 pixels corresponding to approximately 3000 speckles (since Ø=3 pixels). Sampling more speckles can easily be implemented in a typical sCMOS camera with 2048 × 2048 pixels by choosing a larger region of pixels.

As observed in Fig. 6 **b** and **e**, accuracy was seen to be higher at shorter *ρ* and longer *T*, i.e. in the regime of high Φ. Strategies for increasing the amount of detected light to achieve good accuracy while remaining within safety limits may include employing dual sources located equi-distance apart from the detected area of interest.

In addition to Φ(*ρ*), *τ*_*c*_, may also affect accuracy of *k*^2^. In order to study the effect of *τ*_*c*_ on accuracy in *k*^2^, the simulations were repeated fixing Φ (*ρ*) to be constant over all values of simulated *ρ*.

In Fig. 8, the percent error in 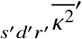 as a function of the number of detected electrons shows that measurement accuracy is dependent on *ρ*, and by extension, *τ*_*c*_. For the simulated camera, measurements with longer *ρ* (shorter *τ*_*c*_) require less detected electrons to achieve the same accuracy in *k*^2^.

**Figure 8.**
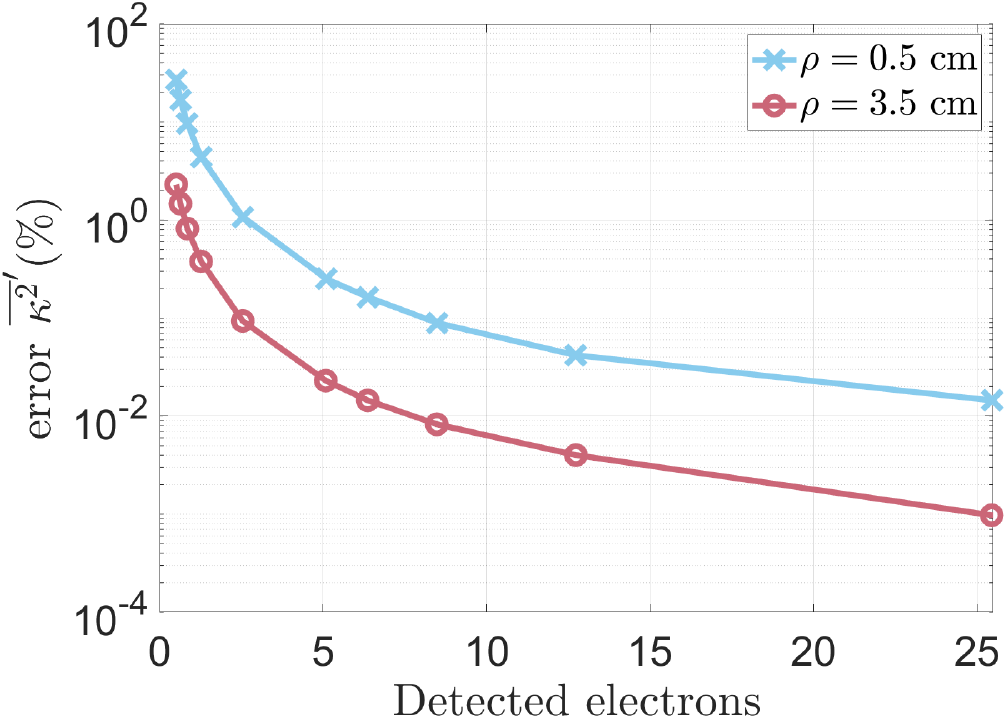
Accuracy of 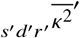 for two different values of *ρ* with identical values of Φ (*T* = 1*ms*). Higher accuracy was found for greater *ρ*.

### 3.5 Using the simulations to design and optimize a system

In the previous sections we have verified the simulation pipeline by comparing the SNR measured experimentally with an Orca Fusion-C14440-20UP camera to the expectations from simulation. We have further demonstrated in detail (without experimental comparison) the entire simulation pipeline. Finally, in the following section we will demonstrate how these simulations can be used to design and optimize a speckle contrast system.

Speckles were simulated using the parameters specified in Table 6. These parameters were derived from the experimental results (*τ*_*c*_ and Ø), properties of the camera defined by the manufacturer, as well as data analysis (*w*_*z*_). The resulting experimental and simulated percent error in *k*^2^ for varying detected electron count rates is shown in Fig. 9.

**Table 6.** Parameters that were used to simulate synthetic speckles based on experimental data taken using a Basler (daA1920-160um) CMOS camera on a liquid phantom.

| Tissue Parameters | Detector Parameters | Speckle Parameters |
| --- | --- | --- |
| $\tau_c : 1.46 \times 10^{-5} \text{ s}$ | QE: 29% | ∅: 5 pixels |
| | $\mu(I_\delta) : 130.9e^-$ | $\epsilon_N : 100$ |
| | $\sigma^2(I_\delta) : 0.8e^-$ | $w_z : [0, 0; 100, 100]$ |
| | $\mu(I_r) : 2.15e^-$ | |
| | $\sigma^2(I_r) : 2.28e^-$ | |

**Figure 9.**
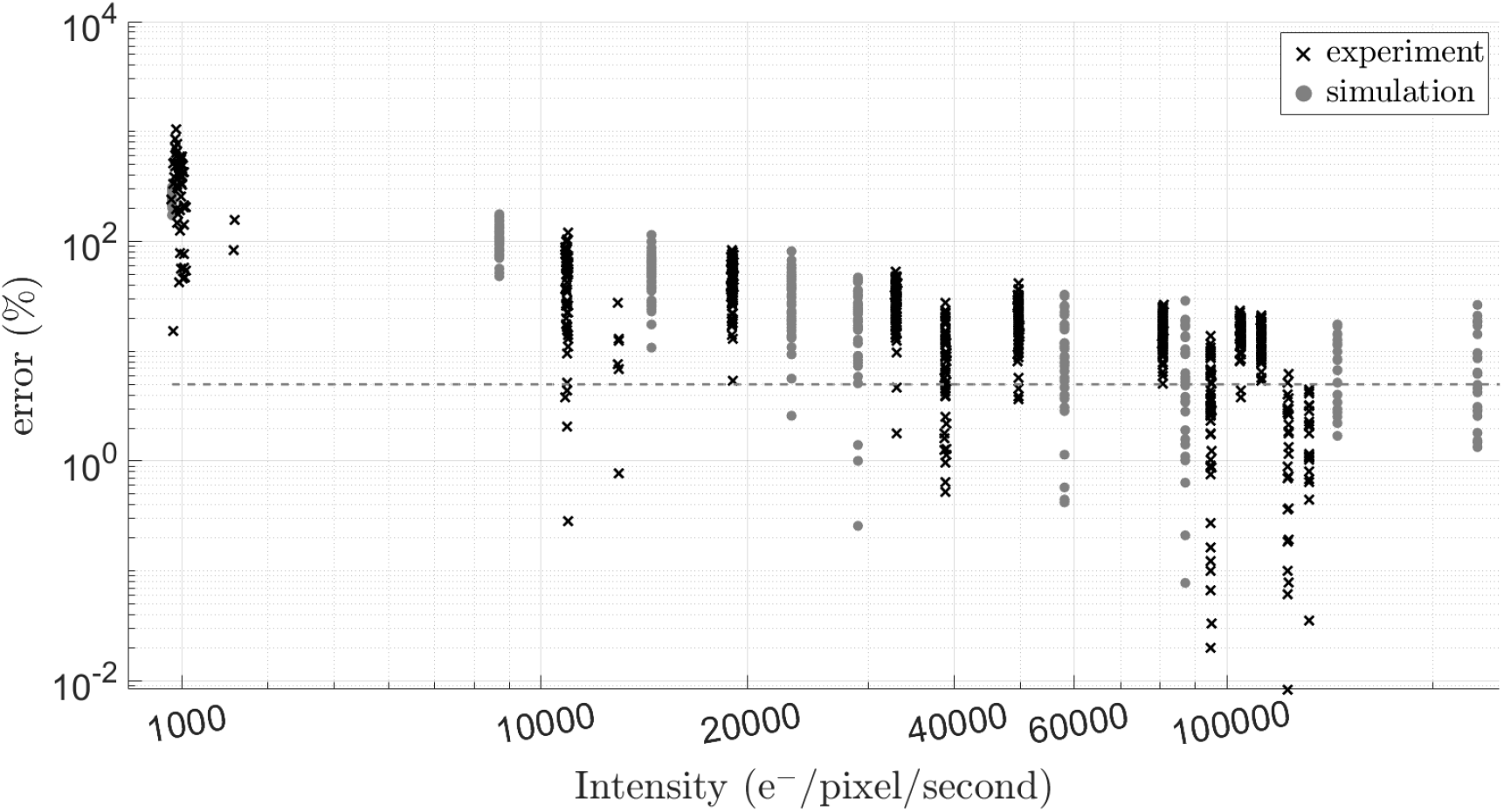
The effect of changing values of detected electron count rate on both the experimental and simulated values of percent error of *k*^2^. The grey horizontal line marks 5% error.

The experimental and simulated results are in good agreement with each other and suggest that for the chosen detector, a minimum detected count rate on the order between 4 to 5 × 10^4^ e^−^/pixel/second allows us to calculate *k*^2^ with approximately 5% error.

Using the derived acceptable minimum detected count rate as a guide in determining the accuracy of raw data signal, the same device was placed on a human subject’s forehead using a *ρ* of 2.53 cm and *T* of 5 ms. Data was acquired at a frame rate of 100 fps. A summary of the measurements is show in Fig. 10. The desired electron count rate was reached (around 4.3 × 10^4^ e^−^/pixel/second, Fig. 10), and the resulting 1 / *k*^2^ shows the expected pulsatile behavior for a measurement acquired at this frame rate (Fig. 10 **a**). In order to confirm that the pulsatile behavior has physiological meaning, the fast Fourier transform (FFT) of the data has also been plotted (Fig. 10 **c**). A distinct peak at 1.4 Hz is seen in the FFT corresponding to a heart rate of 84 bpm. This value matches the resting heart rate measured in this subject using a standard pulse oxymeter. The harmonics of the heart rate can also be seen.

**Figure 10.**
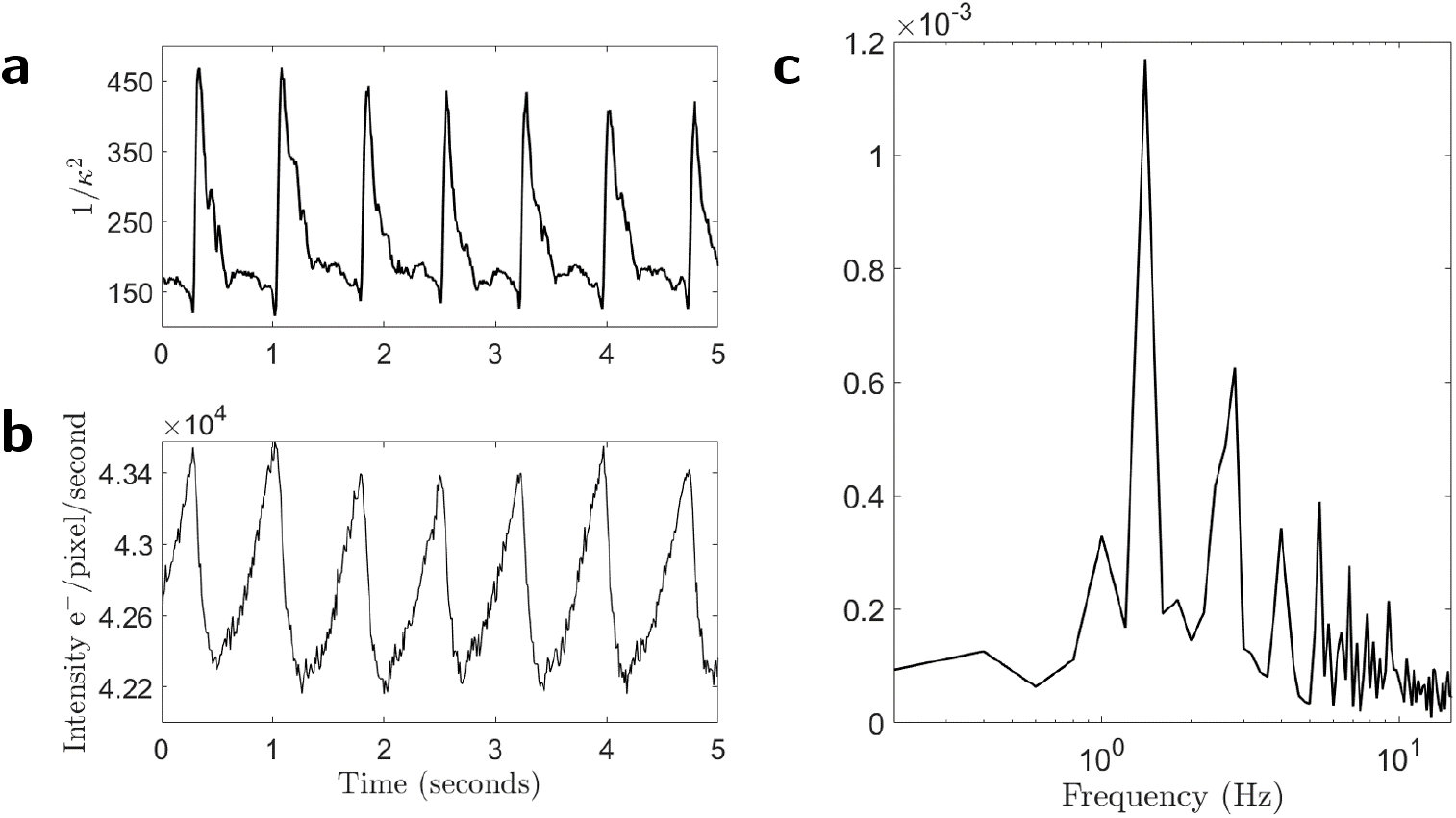
Summary of results from a SCOS measurement on an adult human forehead. 1 / *k*^2^, a surrogate measure of blood flow, shows clear pulsatile signals. **b**) Average detected electron count rate lies in a range which allows us to accurately measure *κ*^2^. **c**) Fourier transform of the *κ*^2^ signal. A clear peak is found at 1.4 Hz corresponding to the heart rate of the subject (84 bpm).

## 4. Discussion

A comprehensive model of speckle contrast signal for measurement of flow requires three main components: the simulation of speckles, their dynamics, and the detector effects on the measured signal. Individual 2D frames of speckles with the correct intensity distribution in these simulations were simulated following the method of Duncan et.al. [55]. The dynamics of the speckle intensity were simulated modifying the method of Ref. [42], where crucially the modification allowed for the characterization of *τ*_*c*_ to be specified according to speckle intensity decorrelation defined by the correlation diffusion equation [10]. While the exact form of the speckle decorrelation, *g*_1_, differs in the simulations, general properties of the dynamics and their dependency on parameters such as *ρ* and *αDb* could be simulated. The simplification of *g*_1_ of a semi-infinite medium as a single exponential function has been seen to be accurate in noise models for DCS [25]. Detector effects were simulated taking into account photon shot noise, dark current signal and noise, and read-out signal and noise. Our method for modeling speckle contrast can account for parameters such as the speckle to pixel size and *β*.

We have shown that the simulations accurately represent experimentally observed behavior of *k*^2^ in the regime of long *ρ* and/or short *T* where the speckle contrast signal increases above the theoretically expected values. Simulation of the noise correction method of Ref. [2] extends the region of *ρ* and *T* where the speckle contrast signal matches its theoretical value. However, depending on the amount of the contribution of the detector effects, the correction cannot account for all of the increased variance from these effects. Therefore, it is important when designing a speckle contrast system to consider the range of *ρ* and *T* where *k*^2^ can be corrected. We have also shown the dependency of accuracy in speckle contrast signal on parameters including the number of detected photons, *ρ*, and *τ*_*c*_.

The accuracy and precision of *k*^2^ developed in the simulation model not only reflects observed experimental behavior, but is also comparable to what has been described in the noise models of related techniques. In DCS, similar to what we have seen in speckle contrast, the SNR of the raw *g*_1_ signal is dependent on the detected photon intensity and *τ*_*c*_. Since DCS uses correlators to measure *g*_1_, the noise model for DCS also depends on the architecture of the correlator [25, 56]. An emerging variation of DCS known as interferometric DCS, or iDCS, utilizes a heterodyne detection technique mixing the traditional DCS signal with a reference arm (i.e. the coherent source). This detection scheme results in greater values of *τ*_*c*_ compared to traditional DCS resulting in an increase in the SNR of the raw *g*_1_ data as well as a decrease in the coefficient of variation of the retrieved blood flow values [15].

While in this analysis we have concentrated on the effects of detector noise in the regime of low detected photon counts corresponding to the typical observations in experiments, it is worth noting that high photon count rates that saturate the detector can also lead to decreases in accuracy as well as precision of the raw signal and in the derived blood flow values. In DCS, saturated detection leads to decreases in the experimentally measured *β* resulting in inaccuracy of the retrieved blood flow [29]. Although not shown here, the same applies in measurements of speckle contrast as detector saturation will lead to inaccurate measurements of *σ*^2^ (*I*) and/or *μ* (*I*) and consequently *k*^2^.

The copula method [55] has previously been used by Qiu et.al. [32] to study the effects of pixel sampling (sampling of *w*_*z*_ and *w*_*t*_) on *k*^2^. In this work, a pseudo exposure time was considered. However since the decorrelation of the speckles were not reassigned in units of time, the simulations were not related to proper physiological or system properties. Thompson et.al [34] combined the method of simulating a single frame of speckles of Ref. [55] with small random phase changes for each consecutively simulated frame, making it very similar to the copula method of Ref. [55]. These simulations were used to study the effect of speckle to pixel size ratio in the measurement of *k*^2^. However, like in Ref. [32], the simulations were not scaled to represent physiological properties and did not include any effects of detector noise.

The present study is complementary to the recent publication by Zilpelwar et.al. [37], with several notable differences. The model developed by Zilpelwar et.al. is based on a Monte-Carlo method simulating random particle (scatterer) motion. Their approach considers a single scattering regime, and is therefore strictly speaking is not applicable for SCOS which is a diffuse optical method considering a multi-scattering regime. Our approach does not simulate particle motion, rather we directly simulate the statistical properties of decorrelating speckle by generating correlated random numbers using the method of Duncan et.al. [42]. Both simulations are based on a single-exponential form of *g*_1_. In the present work, we argue that while the exact value of *k*^2^ is dependent on the approximations used to define *g*_1_, the noise in *k*^2^ is likely not affected due to previous observations in the development of a noise model for DCS [25]. In order to account for the difference in *k*^2^ stemming from discrepancies in the approximation of *g*_1_, in our simulations, we have included a method to correct for this difference. Furthermore, in the present work we were interested in deriving limits of accuracy and precision for an experimental scenario and therefore included a full noise corrected simulation of 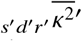 by simulating the expected dark frames of the individual specifications of each simulated camera. These details, multi-scattering regime in a semi-infinite medium, was not included in the model of Ref. [37].

We are not the first to adapt the work of Duncan et.al. [42, 55] to study the behavior of *κ*. We note that this method is not only method in the literature for simulating decorrelating speckle patterns [57–60]. In the copula method of [42], spatial correlation is not preserved between frames. Song et.al propose another method for simulating frames correlated in the spatio-temporal domain [57]. The authors successfully simulated real speckle contrast data by creating correlation maps of data from a rat ear, however the authors note that the accuracy of replicating an image taken from real data depends greatly on the quality of the camera used to acquire the image. Sang et.al. utilized the method of Song et.al. [57] to further expand the method to include time integration effects of exposure time [61], however only one exposure time was simulated. Another method for modelling speckles is to model the summation of random phasors [58]. Postnov et.al. modified this technique in order to simulate the effects of the laser linewidth and camera noise on *k*^2^ [59]. An interesting work by Song et.al. [62] derives the effect of camera quantization of intensity on speckle contrast from the probability density function of speckle intensity. Quantization of the speckle signal is something that was not considered in the current study and should be considered in future work.

## 5. Conclusion

In the present work we have introduced a method for simulating the formation and detection of dynamic speckle patterns. The main application that we have focused on was the design and characterization of a speckle a contrast system capable of measuring human adult cerebral blood flow non-invasively. To this end, the simulation method was validated on a dynamic liquid phantom, the details of speckle contrast signal as a function of *ρ* and *T* were studied, and finally a system designed for human cerebral blood flow was characterized and validated on an adult human subject. The simulation method has been shown to be useful when identifying the lower bounds of detected electron count-rate to achieve the desired accuracy and precision of speckle contrast signal. As speckle contrast signal is sensitive to detector noise effects at low detected electron count-rates, characterizing these limits is advisable when developing any speckle contrast system.

## Data availability

Data underlying the results presented in this paper are not publicly available at this time but may be obtained from the authors upon reasonable request.

